# Genome wide analyses reveal the role of mutator phenotypes in the emergence of drug resistance in *Mycobacterium tuberculosis*

**DOI:** 10.1101/2025.01.17.633521

**Authors:** R. Zein-Eddine, A. Le Meur, S. Skouloubris, L. Jelsbak, G. Refrégier, H. Myllykallio

## Abstract

Antimicrobial combination therapy is widely used to combat *Mycobacterium tuberculosis* (*Mtb*), yet resistance rates continue to rise. Mutator strains, with defects in DNA repair genes, are a known source of resistance in other bacterial infections, but their global role in *Mtb* remains unclear. Here, we provide a comprehensive overview of the contribution of single nucleotide polymorphisms (SNPs) in DNA Repair, Replication, and Recombination (3R) genes to resistance in *Mtb*. Through large-scale bioinformatics analysis of 53,589 whole-genomes, we identified 18 novel SNPs linked to genotypic drug resistance in 3R genes, covering 12.5% of clinical isolates/strains with available genome sequences. Notably, a number of the detected SNPs were positively selected during *Mtb* evolution. Experimental mutation frequency tests further indicated functional defects of these sequence variants in key DNA repair pathways. Our findings indicate that the role of mutability in 3R genes plays a significant role in *Mtb* resistance at the global level.

## Introduction

The social and economic burden of tuberculosis (TB), caused by *Mycobacterium tuberculosis* (*Mtb*), is worsened by the growing issue of antibiotic resistance. Drug resistance in *Mtb* arises exclusively from spontaneous mutations (errors in replication fidelity or nucleotide modifications) usually within chromosomal genes encoding drug targets or enzymes that activate drugs. Unlike many other bacteria, *Mtb* does not acquire resistance through horizontal gene transfer ^1,2,3^. *Mtb*’s genome stability is further threatened by the harsh conditions inside host macrophages, where oxidative stress and antibiotic pressure increase mutation rates ^4,5^. Paradoxically, antibiotics can facilitate the emergence of mutations, including in genes encoding drug targets, driving antimicrobial resistance (AMR) ^6,7^. Mutations in DNA repair genes may give rise to “mutator” phenotypes, which further elevate mutation rates if these genes exhibit reduced function ^8^. Studies in *E. coli* demonstrate that mutator strains can quickly evolve resistance, particularly under combination antibiotic therapy when inhibitory concentrations are delayed. These mutators not only drive multi-drug resistance but may also undermine the effectiveness of combination treatments, favouring resistance under both combination and single-drug regimens ^9^.

Mutator genotypes play a significant role in the emergence of resistance in various bacterial pathogens ^10^. For example, in chronic respiratory infections caused by *Pseudomonas aeruginosa* in cystic fibrosis (CF) patients, mutators frequently emerge due to deficiencies in the DNA mismatch repair (MMR) system, which leads to positive selection for resistant strains^11^. Similarly, high frequencies of mutators have also been observed in CF-related *Staphylococcus aureus* and *Haemophilus influenzae* ^12,13^. In *Mtb*, Previous studies have proposed functional variation in DNA repair genes using limited datasets ^14, 15^,^16, 17^. These studies suggest that *Mtb* strains with altered DNA repair systems may gain fitness advantages under antibiotic pressure. However, they also reveal significant gaps in our current understanding. The existing research is limited in scope, often relying on small datasets (1,600–2,300 isolates) and samples from only a few countries, which may not fully capture the global diversity of *Mtb* strains. Additionally, the impact of mutator genotypes on the development of multidrug resistance in *Mtb* remains insufficiently quantified. Our study aims to address these gaps by investigating a broader and more diverse set of *Mtb* isolates on a global scale, offering new insights into the role of mutator strains in drug resistance.

In this study, we provide a comprehensive analysis of the implications of 3R gene variants (Replication, Recombination, and Repair) in the evolution of molecular resistance mechanisms in *Mtb*. We selected a large panel of 55 genes associated with these mechanisms and explored their diversity using an extensive dataset of clinical and animal isolates from GenBank (54,619 sequence read archive SRA). Through genome-wide association analysis (GWAS), we identified 18 novel variants in different gene systems, all showing high frequencies associated with drug resistance. Additionally, we identified rare variants that emerged independently in various *Mtb* lineages. We predicted the effects of these polymorphisms on protein activity using computational methods and experimentally validated four mutator candidates by assessing their impact on mutation frequency in *Mycobacterium smegmatis* (*Msm*), a model for *Mtb*. This study provides novel insight into our understanding of the role of 3R genes in drug resistance emergence and may contribute to developing accurate diagnostic methods based on whole-genome sequencing (WGS) data for detecting bacteria at risk of developing resistance in clinical settings.

## Results

### Phenotypic drug susceptibility and genotypic profiles of *Mtb* Strains

We analyzed a comprehensive set of 54,619 whole genome sequences, comprising 51,095 clinical *Mtb* strains and 3,524 animal strains sourced from GenBank. Following quality filtering (**see Methods**), the dataset included 53,589 genomes representing *Mtb* strains from various regions worldwide. We developed a high-throughput pipeline to collect SNPs only within a specified list of genes: 1) 55 genes related to the 3R genes, and 2) genes used for strain classification and resistance prediction. All known global *Mtb* lineages infecting humans were represented, including lineage 1 (6.75%), lineage 2 (29.78%), lineage 3 (7.99%), lineage 4 (48.27%), lineage 5 (0.29%), lineage 6 (0.30%), and lineage 7 (0.05%), as well as animal lineages *M. bovis* (6.20%), *M. caprae* (0.22%), and *M. orygis* (0.15%) **(Figure 1b)**.

**Figure 1.**
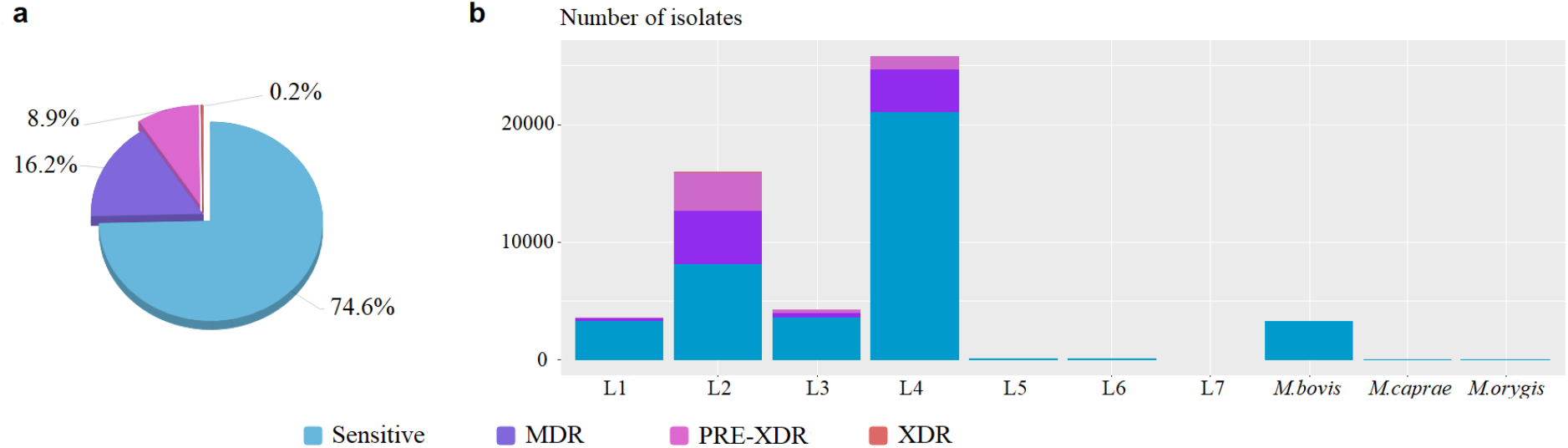
**a.** The pie chart represents the percentage of drug-resistant type of the *Mtb* isolates (n=53,589). **b**. The bar graph represents the proportion of sensitive and drug-resistant strains in each lineage including Sensitive, multidrug-resistant (MDR), pre-extensively drug-resistant (PRE-XDR), and extensively drug-resistant (XDR).

To assess the distribution of drug resistance in this dataset, we computationally predicted resistance for each isolate genome across twenty drugs using the consensus list from TB-Profiler ^18^, a curated compilation of resistance-associated polymorphisms **(see Methods)**. The genotypic analysis of susceptibility to antituberculosis drugs revealed that 39,999 strains (74.7%) were classified as sensitive, 8,689 strains (16.2%) as MDR (carrying mutations conferring resistance to both rifampicin and isoniazid), 4,787 strains (8.9%) as Pre-XDR (MDR genotype with additional resistance to either ofloxacin or kanamycin), and 114 strains (0.2%) as XDR (with mutations for resistance to rifampicin, isoniazid, ofloxacin, and kanamycin) **(Figure 1a)**. In comparison to global resistance rates, we observed the highest resistance rates as expected in lineage 2 and to a lower extent in lineage 4, consistent with findings from other studies ^19, 20^. **(Figure 1b)**.

### GWAS reveals mutator SNPs exhibiting association with drug resistance phenotype in *Mtb*

To identify SNPs contributing to the development of antibiotic resistance in *Mtb*, we conducted an initial screening using allele counting to examine the association between gene adaptive diversity and mutations in 3R genes ^21^. We hypothesized that mutator strains, if they exist, provide a favorable genetic background for the evolution of drug resistance, and are more likely to carry resistance mutations linked to higher mutation rates in MDR, Pre-XDR, and XDR strains. From our SNP-calling pipeline, we generated a comprehensive table (covering 53,589 isolates) that detailed 3R gene mutations, resistance profiles, and classifications for each isolate **(see Methods)**. This dataset was used to statistically assess the relationship between 3R gene SNPs and drug resistance through chi-square tests and Fisher’s exact tests for smaller SNP samples. This approach aimed for high sensitivity, acknowledging the typical trade-off with specificity found in allele-counting methods ^21^. These tests, without accounting for population structure and recombination, identified 156 missense SNPs across 43 out of 55 3R genes, potentially linked to antibiotic resistance (p-value < 0.05) **(Supplementary Table S1)**.

We then performed a second screening using TreeWAS ^22^ to improve specificity by filtering out false positives from the initial analysis. TreeWAS is a tool for Genome-Wide Association Studies that considers population structure and recombination, making it particularly useful for clonal organisms like *Mtb*. Incorporating evolutionary relationships improves the accuracy of detecting true genetic associations with phenotypes, such as drug resistance. TreeWAS calculates three scores—terminal, simultaneous, and subsequent—using phenotype and genotype trees. The terminal score correlates leaf phenotypes and genotypes, the simultaneous score tracks parallel changes in branches, and the subsequent score assesses the proportion of branches with concurrent phenotype-genotype changes. We applied TreeWAS to each 3R gene individually to identify associations with drug resistance profiles.

Overall TreeWAS identified 18 non-synonymous SNPs in 3R genes significantly associated with resistance (using the terminal and subsequent scores, p < 0.000001). All were part of the 156 SNPs detected using the allele counting method **(Table 1)**. Among these, five SNPs were supported by the three scores and affected *xthA* **(Figure 2a.b)**, *mutT3, recG*_*wed*_, *recN*, and *uvrD2* **(Table 1)**. These 18 SNPs were found in lineage 2 and/or in lineage 4, with frequencies ranging from 0.05% (Trp196Arg in MutT3) to 1.99% (Glu331Gly in UvrD2) altogether concerning 6,650 clinical samples (**Table 1)**. They are considered candidate mutators in the subsequent sections.

**Table 1.**
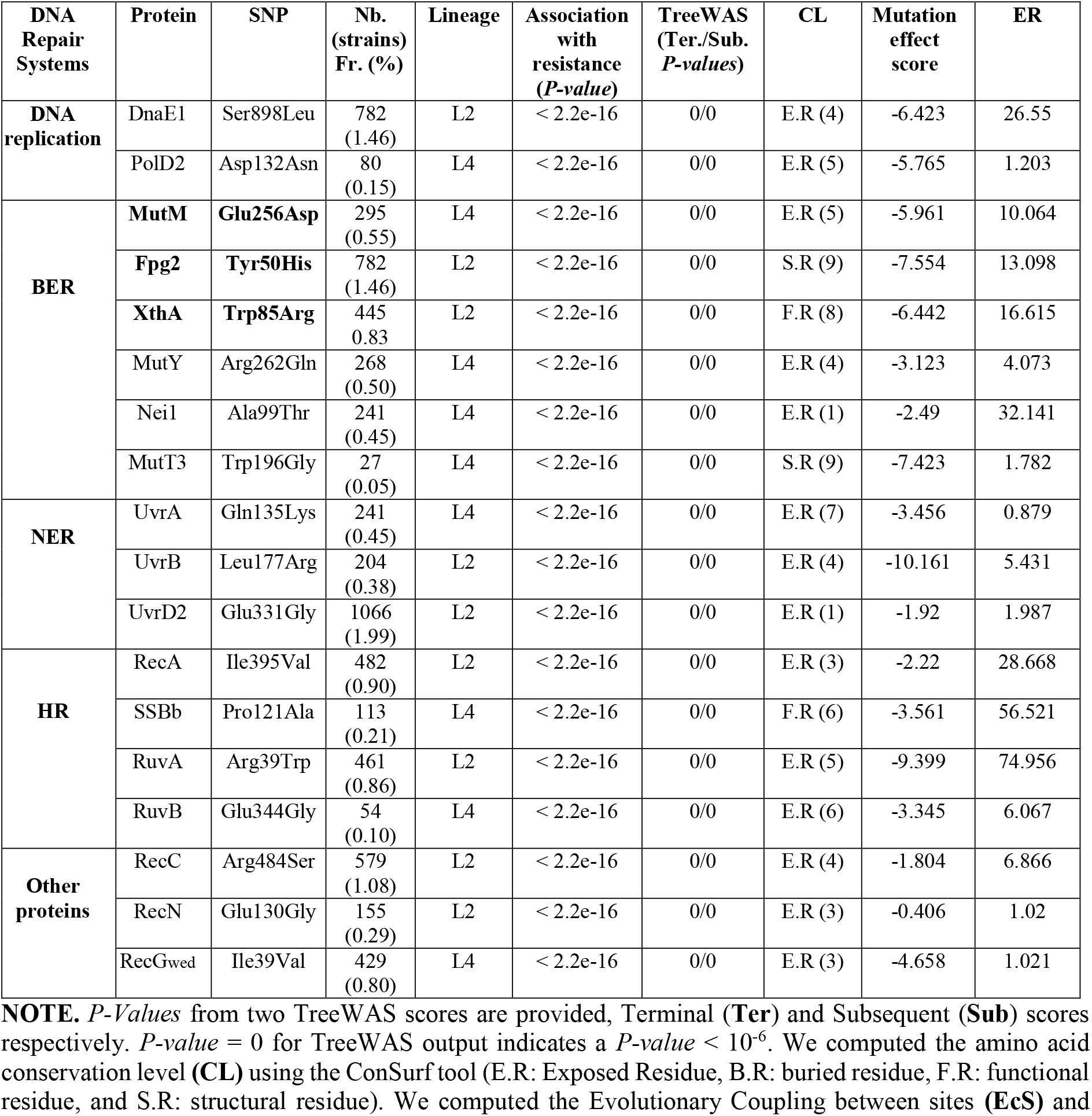
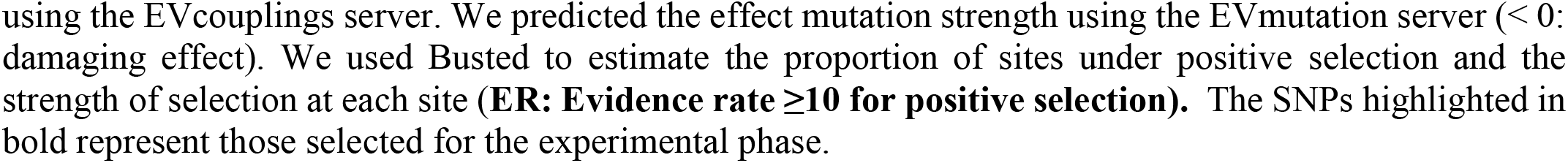
GWAS and computational analyses results of SNPs associated with drug resistance phenotype.

| DNA Repair Systems | Protein | SNP | Nb. (strains) Fr. (%) | Lineage | Association with resistance (*P-value*) | TreeWAS (Ter./Sub. *P-values*) | CL | Mutation effect score | ER |
| --- | --- | --- | --- | --- | --- | --- | --- | --- | --- |
| DNA replication | DnaE1 | Ser898Leu | 782 (1.46) | L2 | < 2.2e-16 | 0/0 | E.R (4) | -6.423 | 26.55 |
|  | PolD2 | Asp132Asn | 80 (0.15) | L4 | < 2.2e-16 | 0/0 | E.R (5) | -5.765 | 1.203 |
| BER | MutM | Glu256Asp | 295 (0.55) | L4 | < 2.2e-16 | 0/0 | E.R (5) | -5.961 | 10.064 |
|  | Fpg2 | Tyr50His | 782 (1.46) | L2 | < 2.2e-16 | 0/0 | S.R (9) | -7.554 | 13.098 |
|  | XthA | Trp85Arg | 445 0.83 | L2 | < 2.2e-16 | 0/0 | F.R (8) | -6.442 | 16.615 |
|  | MutY | Arg262Gln | 268 (0.50) | L4 | < 2.2e-16 | 0/0 | E.R (4) | -3.123 | 4.073 |
|  | Nei1 | Ala99Thr | 241 (0.45) | L4 | < 2.2e-16 | 0/0 | E.R (1) | -2.49 | 32.141 |
|  | MutT3 | Trp196Gly | 27 (0.05) | L4 | < 2.2e-16 | 0/0 | S.R (9) | -7.423 | 1.782 |
| NER | UvrA | Gln135Lys | 241 (0.45) | L4 | < 2.2e-16 | 0/0 | E.R (7) | -3.456 | 0.879 |
|  | UvrB | Leu177Arg | 204 (0.38) | L2 | < 2.2e-16 | 0/0 | E.R (4) | -10.161 | 5.431 |
|  | UvrD2 | Glu331Gly | 1066 (1.99) | L2 | < 2.2e-16 | 0/0 | E.R (1) | -1.92 | 1.987 |
| HR | RecA | Ile395Val | 482 (0.90) | L2 | < 2.2e-16 | 0/0 | E.R (3) | -2.22 | 28.668 |
|  | SSBb | Pro121Ala | 113 (0.21) | L4 | < 2.2e-16 | 0/0 | F.R (6) | -3.561 | 56.521 |
|  | RuvA | Arg39Trp | 461 (0.86) | L2 | < 2.2e-16 | 0/0 | E.R (5) | -9.399 | 74.956 |
|  | RuvB | Glu344Gly | 54 (0.10) | L4 | < 2.2e-16 | 0/0 | E.R (6) | -3.345 | 6.067 |
| Other proteins | RecC | Arg484Ser | 579 (1.08) | L2 | < 2.2e-16 | 0/0 | E.R (4) | -1.804 | 6.866 |
|  | RecN | Glu130Gly | 155 (0.29) | L2 | < 2.2e-16 | 0/0 | E.R (3) | -0.406 | 1.02 |
|  | RecGwed | Ile39Val | 429 (0.80) | L4 | < 2.2e-16 | 0/0 | E.R (3) | -4.658 | 1.021 |

**NOTE.** *P-Values* from two TreeWAS scores are provided, Terminal (**Ter**) and Subsequent (**Sub**) scores respectively. *P-value* = 0 for TreeWAS output indicates a *P-value* < 10<sup>-6</sup>. We computed the amino acid conservation level (**CL**) using the ConSurf tool (E.R: Exposed Residue, B.R: buried residue, F.R: functional residue, and S.R: structural residue). We computed the Evolutionary Coupling between sites (**EcS**) and
using the EVcouplings server. We predicted the effect mutation strength using the EVmutation server ( $< 0$ : damaging effect). We used Busted to estimate the proportion of sites under positive selection and the strength of selection at each site (**ER: Evidence rate $\geq 10$ for positive selection**). The SNPs highlighted in bold represent those selected for the experimental phase.

**Figure 2.**
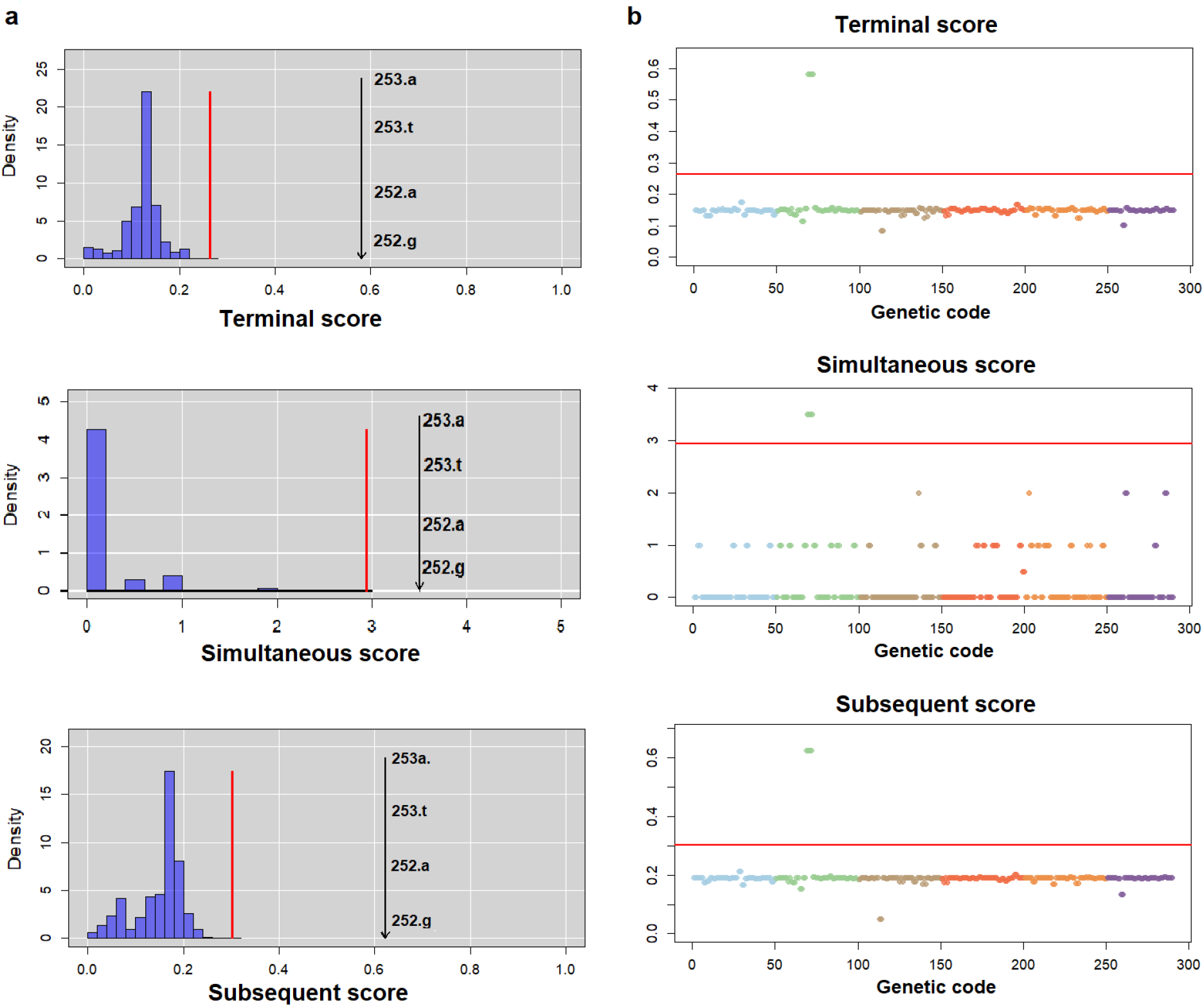
The TreeWAS results identify the Trp85Arg SNP in the *xthA* gene as being significantly associated with antibiotic resistance. **a.** The null distributions of simulated associations for the three TreeWAS scores-terminal, simultaneous, and subsequent are shown. The Trp85Arg SNP in *xthA* (nucleotide position/gene: 252 and 253) is identified above the threshold in all three score distributions, indicating a significant association. **b**. The Manhattan plots visualize SNP associations (amino acid position/gene), with the red line marking the significance threshold. SNPs surpassing this threshold, including the Trp85Arg variant, are highlighted based on their scores. The colours of the data points were randomly assigned for differentiation across the three scores.

To further investigate the distribution of these 18 SNPs across the different 3R genes, we conducted a principal component analysis (PCA) to determine if significance clustered according to specific functions among the 3R genes. This analysis did not reveal a distinct system-specific cluster **(Supplementary Figure S1)**, suggesting that all DNA repair and replication systems may contribute to the emergence of mutator genotypes associated with increased resistance.

### Positive selection and damaging effects of SNPs associated with drug resistance

To investigate how selection has influenced the evolution of 3R genes, we conducted branch-site tests using BUSTED ^23^. This type of analysis detects episodic diversifying selection and assesses individual codons likely under positive selection within each gene. Among the 18 genes analyzed, 11 exhibited significant signatures of positive selection (FDR-adjusted p-values < 0.05), including *dnaE1, mutM, xthA, nei1, mutT3, uvrA, uvrB, recA, ssbB*, and *recB* **(Supplementary Table S2)**. Notably, 8 of the 18 SNPs identified by TreeWAS as being associated with drug resistance were under diversifying positive selection **(Table 1, ER)**.

To evaluate the potential phenotypic damage caused by these SNPs, we analyzed their ability to disrupt protein function, structure, stability, or expression *in vivo* using the EVmutation tool ^24^. This tool models epistasis by accounting for interactions between protein residues and predicts the deleterious effects of mutations. We selected EVmutation because it provides precomputed predictions for all possible amino acid substitutions in most mycobacterial DNA repair proteins, classifying mutations as beneficial, neutral, or damaging. Our analysis revealed that all 18 SNPs associated with drug resistance were classified as damaging, with effect scores ranging from -2.49 to -10.16, indicating highly damaging effects **(Table 1). Phylogenetic features of candidate mutator SNP isolates**

To investigate the evolutionary dynamics of *Mtb* strains carrying candidate mutator SNPs, we retrieved whole-genome SNPs distinguishing two sets of *Mtb* isolates within the same sub-lineage: those carrying the target mutator candidate and those without it. SNPs in repetitive regions were excluded from the analysis. We constructed maximum likelihood (ML) phylogenies using the GTR+Γ+I substitution model. The GTR is built on the molecular diversity present in the samples and is widely used in *Mtb* evolutionary studies. To further refine the model, gamma-distributed rate variation (+Γ) was incorporated to capture evolutionary rate heterogeneity, while invariant sites (+I) accounted for conserved regions.

Specifically, we analyzed 84 whole genomes, including 39 isolates with the Trp85Arg SNP in XthA and 45 control isolates from the same sub-lineage (L2.2.1), which lacked this SNP or any other resistance-associated mutations in the 3R genes. The majority of isolates carrying the Trp85Arg SNP exhibited multidrug-resistant (MDR) or pre-extensively drug-resistant (PRE-XDR) phenotypes (**Figure 3a**). Phylogenetic analysis revealed that isolates with this SNP clustered in a distinct branch with strong bootstrap support (100%). This cluster is characterized by an unexpectedly long branch compared to other groups of isolates **(Figure 3b)**. Similar analyses were conducted for the other candidate SNPs when control group strains of the same sub-lineage were available, including those in d*naE1, polD2, mutM, uvrA, mutY, nei1, uvrD2, recA, recC*, and *recN*. In these reconstructions, long branches were consistently observed, with a few terminal branches also being relatively long **(Supplementary Figures S2)**.

**Figure 3.**
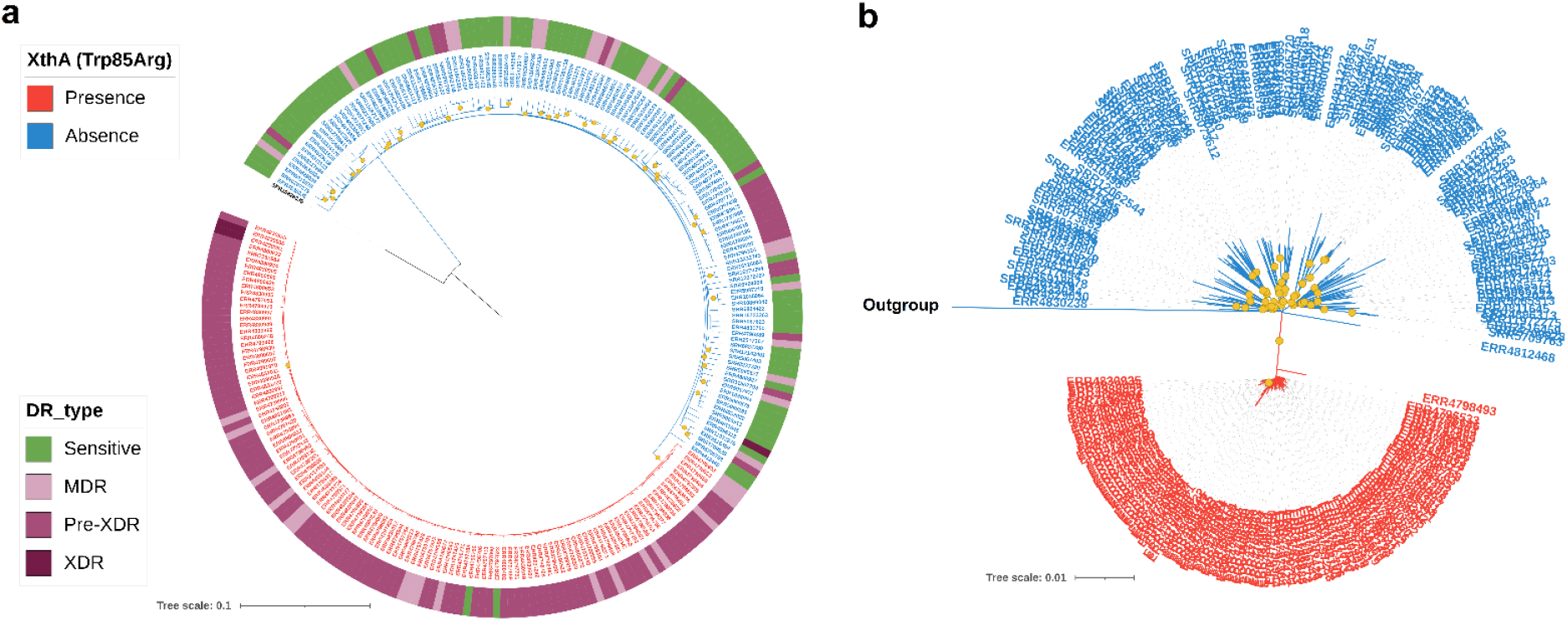
Maximum Likelihood (ML) Tree of *Mtb* Genomes. **a.** ML tree of 85 *Mtb* genomes, illustrating the presence or absence of the XthA SNP (Trp85Arg) and the associated drug-resistant profiles of the isolates. **b**. An unrooted form of the ML tree illustrating the presence or absence of the XthA SNP (Trp85Arg) and showing the evolutionary relationships among these strains. The tree is drawn to scale, with branch lengths representing the number of substitutions. The isolate SRR10828835 (lineage 8) was used as an outgroup for all analyses. Bootstrap support values of 100% (based on 1000 replicates) are marked by yellow dots on the internal branches. The branch and label colours represent the presence of the SNP (red) and the absence of the SNP (blue). The colour strip indicates the drug-resistant profiles (Sensitive, MDR, PRE-XDR, and XDR) of the isolates.

To verify the accuracy of these long branches, we performed pairwise distance tests, comparing genetic distances (quantified by sequence divergence) to tree-based distances. These tests confirmed that the branch lengths reflect true evolutionary distances rather than being artefacts of the reconstructions. The results of the Mantel tests further supported this conclusion, with a Mantel statistic of 0.99 and a *p-value* of 0.001, indicating a highly significant correlation between the genetic distances and tree-based distances. Pearson and Spearman correlation coefficients were both 0.99, demonstrating a very strong linear and monotonic relationship between these two distance measures.

### Mutator effect for the selected SNPs in the non-pathogenic *Msm* model

Our next objective was to assess the impact of consistent candidate mutators on mutation frequency in vitro. In addition to the analyses performed earlier (BUSTED and EVmutation), we estimated the evolutionary conservation of each amino acid position to predict the mutational impacts on the structure and function of the corresponding protein using the ConSurf tool ^25^ **(Supplementary Table 1)**. We focused on highly conserved SNP positions that demonstrated evidence of positive selective pressure at the codon site, with damaging mutation effect strengths of ≤ -5. Additionally, the selected SNP positions needed to be conserved in *Msm*, as this organism was utilized as a model species for *Mtb*, consistent with previous mutagenesis studies. Out of the 18 SNPs identified as associated with drug resistance using TreeWAS, three candidates met all the above criteria: MutM (Glu256Asp), Fpg2 (Tyr50His), and XthA (Trp85Arg) (marked in bold in **Table 1**). The frequencies of these three alleles in the studied population ranged from 0.55% for Glu256Asp to 1.46% for Tyr50His **(Table 1)**.

In addition to the three selected SNPs, we aimed to investigate a mutation in *nucS*, a novel gene involved in the mismatch repair (MMR) system in *Mtb*. NucS corrects mismatched bases that occur during DNA replication, thereby preventing point mutations ^26, 27^. We focused on the Tyr132Ser mutation in NucS, which was not detected by TreeWAS due to its low frequency in the studied population (0.01%). However, this mutation was identified in two independent lineages (L2 and L3), indicating a rare instance of convergence **(Supplementary Table S1)**. Furthermore, the codon position for this mutation exhibited strong selection (40.73) during evolution, as determined by the BUSTED analysis **(Supplementary Table S1)**, highlighting a functional nuclease domain in *nucS* with a damaging effect score of -5.161.

To test the effects of the selected four allelic polymorphisms on the activity of different genes, we constructed three mutant alleles by introducing point mutations at the appropriate positions in the chromosome of *Msm* through allelic exchange. For the NucS (Tyr132Ser) allele, we employed the CRISPR-Cas12a-assisted recombineering technique (see Methods). We assessed the phenotypes of the constructed strains by measuring spontaneous resistance to rifampicin and isoniazid. Hyper-mutability was defined as a 10-fold increase in mutation frequency compared to the wild-type allele **(Figure 4a)**. As anticipated, the four alleles (MutM (Glu256Asp), Fpg2 (Tyr50His), XthA (Trp85Arg), and NucS (Tyr132Ser)) resulted in increases in mutation frequency of 10, 14, 14, and 18 folds, respectively, for rifampicin, and 20, 18, 19, and 24 folds, respectively, for isoniazid **(Figure 4b)**. These experimental results suggest that these four SNPs cause DNA repair defects in Mycobacteria.

**Figure 4.**
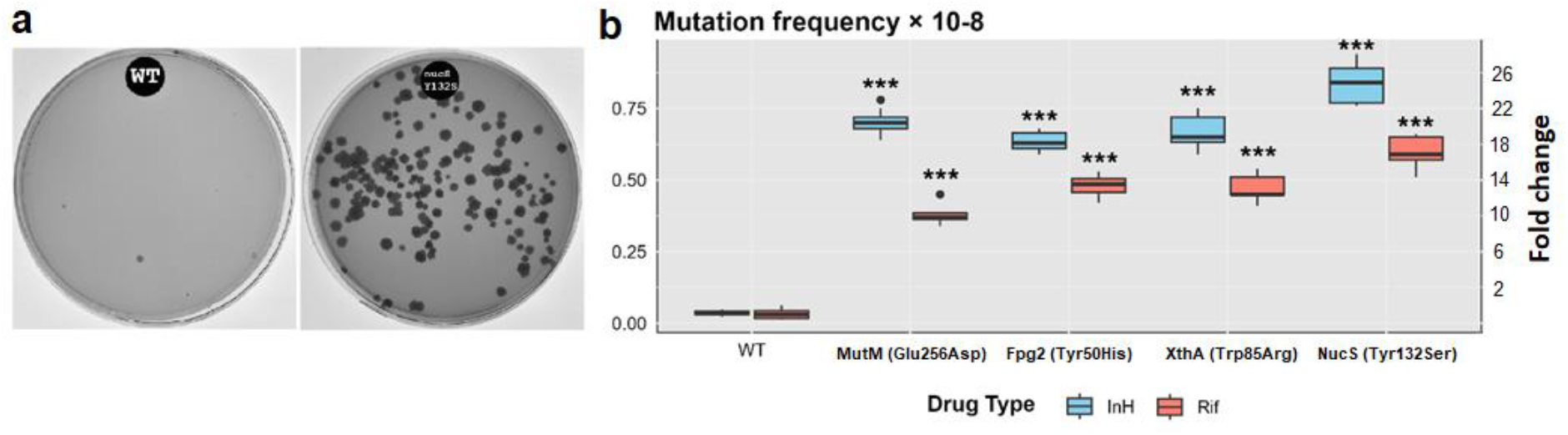
Experimentally Determined Mutational Effects of Different Mutator Variants. **a.** Frequency of spontaneous Rifampicin-resistant mutants compared to that of the wild-type (WT), demonstrating the hypermutable phenotype. **b**. Frequency of spontaneous mutations conferring resistance to rifampicin and isoniazid in *Mycobacterium smegmatis* mc2155 (WT) and the various mutants. The box plot also displays the fold-change in mutation frequencies of the different mutants compared to the wild type of *Msm*. Three biologically independent sets of experiments were conducted, with each biological experiment performed in triplicate. Statistical analysis (one-way ANOVA) was conducted using the R base package, with a significance level indicated as ***p < 0.0001.

## Discussion

The global rise in *Mtb* antimicrobial resistance demands innovative and measurable strategies to stop its spread. Effective approaches, such as combination antibiotic therapy, are urgently needed. This strategy uses multiple drugs simultaneously, making resistance rare, as it requires several mutations to occur at once in the same genetic background ^28^. Under these conditions, wild-type bacteria generally cannot develop resistance. However, mutator strains with defects in the 3R genes can more rapidly evolve multi-drug resistance ^9^. Thus, early identification of these mutator strains is crucial for enhancing the effectiveness of combination therapies and improving treatment outcomes. In this study, we first applied a bioinformatics strategy using whole-genome sequencing data to extract mutations in 3R and antibiotic-resistance genes. Next, we conducted a GWAS to identify potential mutator candidates associated with genotypic drug resistance, revealing further evidence of their mutator nature. Computational tools were then used to prioritize promising candidates, which were experimentally validated by introducing point mutations into the *Msm* chromosome, followed by mutation frequency tests.

Our dataset, comprising 53,589 SRAs, represents the global diversity of natural *Mtb* isolates. This extensive data enabled us to identify 18 SNPs associated with drug resistance across 18 genes involved in various DNA repair pathways **(Table 1)**. Previous studies showed that mutators in clinical populations mainly arise from defects in the MMR pathway, leading to significant mutations that enhance bacterial survival ^29,30, 31^. These studies typically highlight mutations in key MMR genes such as *mutS, mutL*, and *mutH* as the primary drivers of mutator phenotypes, with *mutS* mutations being the most common ^32^.

However, our findings suggest that in natural *Mtb* isolates, mutators do not arise from defects in the MMR pathway. Instead, they are linked to mutations in other DNA repair genes, suggesting that the mutator phenotype in *Mtb* involves multiple DNA repair pathways. This highlights a more complex interplay of DNA repair systems during *Mtb* infections. The harsh chemical environment faced by *Mtb* likely causes diverse forms of DNA damage, beyond just replication errors, reinforcing the role of various repair pathways in shaping the mutator phenotype.

Previous studies have identified three missense SNPs in the proofreading domain (PHP) of the DNA polymerase *dnaE1*^14^ and five SNPs in the MMR component *nucS* ^16^ linked to mutator phenotypes. However, none have established an association between these SNPs and drug resistance profiles in natural *Mtb* isolates. This lack of association may be attributed to their rare occurrence in the population. In addition, our analysis identified two SNPs (MutY_Arg262Gln and UvrA_Gln135Lys) **(Supplementary table S1)**, previously reported as associated with drug resistance ^17^, but these two SNPs did not show significant association with genotypic resistance in our study. This difference may be due to their reduced impact, dragged down in our much larger dataset (53,589 SRA vs. 2,773 SRA), or due to reduced opportunities for the strains carrying these mutations to acquire resistance because of their emergence in a context of tighter surveillance or reduced use of antibiotics. In contrast, we identified new SNPs likely because of our increased statistical power due to our large sample size. Moreover, we focused on genotypic resistance and not phenotypic resistance. We further validated the hyper-mutability of four SNPs, demonstrating increases in mutation frequency exceeding tenfold in our experimental work **(Figure 4b)**. Hyper-mutators in clinical isolates often exhibit 10 to 100-fold increased mutation rates ^33, 34^. These elevated mutation rates can facilitate the selection of hyper-mutators, promoting the progression of chronic diseases and driving the evolution of resistance to therapeutic agents ^34^. Also, our findings indicate that these SNPs have undergone positive selection during the evolution of *Mtb*, which strengthens our finding and indicates a link between hyper-mutability and antibiotic resistance in *Mtb* patients.

All 18 SNPs associated with resistance identified through GWAS were located in lineage 2 (9 SNPs) and lineage 4 (9 SNPs) **(Table 1)**. Our study, along with previous research, has shown that *Mtb* strains from lineage 2 and lineage 4 are more likely to develop resistance **(Figure 1b)** and maintain fitness in the presence of resistance-conferring mutations compared to other lineages ^35,36,37^. Here, we hypothesize that one possible explanation for the emergence of resistance in these two lineages could be linked to the presence of mutator SNPs, which increase mutation rates. These mutator SNPs may create a favorable environment for adaptive mutations to occur more frequently, accelerating the evolution of resistance while preserving the overall fitness of the bacterial population.

Our analysis showed a significant prevalence of candidate mutators in *Mtb*, with 12.5% of the isolates carrying a consistent candidate mutator SNP **(Table 1)**. This suggests that mutator candidates are surprisingly common within the *Mtb* population. Interestingly, not all candidate mutator strains exhibited genotypic resistance. A possible explanation for this is that SNPs in DNA repair pathway genes may promote the accumulation of other beneficial mutations in bacteria under stress, beyond those directly related to drug resistance. A subset of these mutations may indeed facilitate the organism’s adaptation to various selection pressures, such as the host immune response, while also enabling the bacteria to acquire resistance to multiple antibiotics sequentially or simultaneously. Of note, the increased genetic mutation rates may enhance the potential for developing resistance to second-line drugs even during a period with first-line treatment ^9^.

The phylogenetic analysis revealed that, for each candidate mutator SNP, strains harboring the SNP clustered together in a single branch node, which was highly associated with drug resistance profiles **(Figure 3a)** and characterized by a long branch compared to other groups of isolates **(Figure 3b)**. This pattern indicates a greater genetic divergence for these clades, suggesting that the corresponding strains have accumulated more mutations over a certain period. However, the terminal branches were mostly short. We propose that the introduction of these SNPs in the DNA repair genes may have initially contributed to an increased accumulation of mutations. The persistence of these mutations was then influenced by specific selective pressures, either related to treatment, host defense, or both. In a subsequent phase, the mutation rate may have dropped due to compensatory mutations. Hypermutable phenotypes can indeed confer advantages, such as increased adaptability, but they also come with the cost of accumulating deleterious changes. Once a mutator strain acquires an adaptive trait, such as drug resistance, it is expected to evolve back towards a lower mutation rate ^38, 39^. We speculate that the candidate mutator strains detected may currently exhibit lower mutation rates than they did in their recent history due to compensatory mutations elsewhere in the genome. Furthermore, the hyper-mutability observed in *Msm* may exceed that in *Mtb* due to the complex interaction of 3R genes. Further experimental investigation of clinical strains or *Mtb*-engineered strains carrying these SNPs is needed to better understand 3R gene interactions. Additionally, the variations in these strains, which likely provide advantageous traits, may offer insights into *Mtb* pathogenesis, including virulence, transmissibility, and persistence in TB patients.

In conclusion, our study provides a detailed genome-wide catalogue of SNPs in 3R genes linked to drug resistance in natural *Mtb* isolates, offering the potential for use in drug resistance screening to inform treatment strategies. This research enhances our understanding of the genetic basis of drug resistance and highlights the potential of monitoring these SNPs in clinical settings to improve resistance management and patient outcomes.

## Methods

### Characterization of *Mtb* diversity in DNA repair genes

A pipeline was developed and optimized to download and identify SNPs and small insertions and deletions (indels) in regions corresponding to an ancestral version of *Mtb* based on the H37Rv reference genome ^40^, with parameters favoring sensitivity. Briefly, sequence reads were retrieved from the European Nucleotide Archive (ENA) (https://www.ebi.ac.uk/ena/) and the NCBI database (https://www.ncbi.nlm.nih.gov/) using the SRA toolkit. For each sample, Fastp software ^41^ was utilized for adapter trimming and read filtering. Nucleotide positions in the reads with a quality score below Q25 were discarded. Quality profiling was checked with FastQC ^42^. High-quality reads were then aligned to the ancestral reference genome using BWA-MEM2 software ^43^, which is based on the H37Rv sequence and shares identical genome annotations ^40^. Single nucleotide polymorphisms (SNPs) and small indels were called using the Genome Analysis Toolkit (GATK) Haplotype caller v4 ^44^. Sample genotypes were determined by selecting the majority allele according to default parameters, with a minimum quality threshold of 20. Positions with insufficient support were classified as missing data. This approach prioritizes sensitivity at the expense of specificity. Specificity is ensured through the independent identification of SNPs across multiple samples. To optimize data quality and storage, SNPs were then extracted from a predefined list of genes of interest, which included 55 genes from various 3R pathways, as well as genes important for classifying the isolate and predicting its resistance profile (see below).

### Identification of lineage and drugs resistant profile *in silico*

The pipeline was also utilized for two main objectives: (1) to identify all mutations associated with antimicrobial resistance (AMR) according to the catalogue of mutations in *Mtb* published by the WHO in 2022 *i*.*e*. to obtain genotypic resistance, and (2) to identify classification SNPs as outlined in previous studies ^45,46,47,48^. Specifically, each sample was characterized as resistant or susceptible to common drug types, categorized as follows: fully sensitive (S), multidrug-resistant (MDR), pre-extensively drug-resistant (Pre-XDR), and extensively drug-resistant (XDR). Additionally, samples were assigned to known tuberculosis lineages or sub-lineages. A total of twenty-one drugs were included in the genome-wide analysis, comprising isoniazid (INH), rifampicin (RIF), ethionamide (ETH), pyrazinamide (PZA), ethambutol (EMB), streptomycin (STM), amikacin (AMK), capreomycin (CAP), kanamycin (KAN), ciprofloxacin (CIP), ofloxacin (OFL), moxifloxacin (MOX), cycloserine (CYS), and para-aminosalicylic acid (PAS). Drug family groups, including second-line injectable drugs (SLIDs: AMK, KAN, and CAP) and fluoroquinolones (FLQs: CIP, OFL, and MOX), were also analyzed. Samples with ambiguous resistance or classification outputs were excluded from further analysis.

### Allele counting association analysis

A summary table of mutations and isolates was created to conduct chi-square or Fisher’s exact tests, depending on the sample sizes, to identify significant associations between the presence of mutations in 3R genes and drug resistance. All analyses were performed using the R base package ^49^.

### GWAS using a phylogenetic tree-based approach

A phylogenetic method for conducting Genome-Wide Association Studies (GWAS) was implemented using the TreeWAS package in R ^22^. TreeWAS assesses the statistical association between the drug-resistant profiles of isolates and their genotypes at all loci, identifying significant associations while accounting for the confounding effects of clonal population structure and homologous recombination. This method also computes the homoplasy distribution, which includes site-specific substitution counts derived from the empirical dataset using the Fitch parsimony algorithm. Association testing between each genetic locus and phenotype is performed through three independent tests for each locus ^22^. We conducted TreeWAS analyses on each of the 43 genes individually, looking for associations between variants within the 3R genes and genotypic drug resistance profiles. Concatenated SNP alignments over the whole genome were generated using TB Annotator ^50^. This platform allows easy selection of samples, implements the GATK variant caller, and ensures stringent selection of SNPs in non-repetitive regions. The SNP alignments were utilized to reconstruct a maximum-likelihood phylogeny with FastTree v2.1.8 ^51^. Subsequently, this phylogeny, along with the resistance profile matrix (sensitive/resistant) and the multiple sequence alignment for each of the 3R genes, served as inputs for TreeWAS v1.0 ^22^.

### Selection Analysis

Positive selection analysis was conducted using the Branch-site Unrestricted Statistical Test for Episodic Selection (BUSTED) ^23^. This analysis aimed to (1) assess each of the 3R genes for evidence of episodic selection utilizing a pre-calculated phylogenetic tree, and (2) compare the rates of synonymous (silent) and non-synonymous (amino acid-changing) substitutions at each codon site.

### Compute mutation effects strength predicted from sequence co-variation

The EVmutation server ^24^, which employs evolutionary couplings from sequence covariation, was utilized with default parameters to calculate the quantitative effects of each mutator candidate on the stability and function of the corresponding protein.

### Estimate the evolutionary conservation of amino acid positions

The ConSurf web server ^25^ was employed with default parameters (E-value cutoff <0.0001) to assess the evolutionary conservation of amino acid positions in the 3R proteins of interest. This analysis utilized a probabilistic framework grounded in the phylogenetic relationships among homologous sequences, following a Bayesian method.

### Principal component analysis

Principal Component Analysis (PCA) was conducted using the stats package in R ^49^. The analysis was performed with data centering but without scaling. Subsequently, the PCA results were visualized using the ggplot2 package ^52^.

### Phylogenetic trees construction and pairwise-distance analyses

To reconstruct the phylogenetic trees, a multiple sequence alignment of concatenated SNPs from the whole *Mtb* genome was obtained using TB-annotator ^50^. We inferred maximum likelihood (ML) phylogenies for two sets of *Mtb* isolates of the same sub-lineage—those carrying and lacking the target mutator candidate— using RAxML-HPC ^53^ with the GTR+Γ+I substitution model. The GTR is built on the molecular diversity present in the samples and is widely used in *Mtb* evolutionary studies. The gamma-distributed rate variation (+Γ) captures evolutionary rate heterogeneity, while invariant sites (+I) address conserved regions. Mutations in repetitive areas were excluded from the analysis. The isolate SRR10828835 (lineage 8) served as an outgroup for all analyses. A total of 1,000 bootstrapped datasets were utilized to estimate the statistical confidence of the nodes, and ascertainment bias associated with using only polymorphic sites was corrected using Felsenstein’s correction in RAxML. The resulting phylogenetic trees were visualized using the iTol tool software ^54^.

The pairwise distances from the aligned FASTA sequences were calculated using the ape package in RStudio ^55^, which employs the Jukes-Cantor model to assess genetic divergence. A phylogenetic tree was constructed using RAxML as described earlier and then downloaded to RStudio for further analysis. We performed correlation tests between the tree distances and pairwise genetic distances, calculating both Pearson and Spearman correlation coefficients for robustness. Additionally, a Mantel test was conducted using the vegan package ^56^ in RStudio to assess the correlation between these distance matrices, yielding significant results. Finally, relationships were visualized using ggplot2, with scatter plots and linear regression lines constructed to illustrate the correlations.

### Strains, media, and growth conditions

*Escherichia coli (E. coli)* strains DH5α and XL10, along with *Mycobacterium smegmatis* (*Msm*) mc^2^155 (GenBank: CP000480), were utilized in this study. *E. coli* strains were cultured at 37°C in LB medium supplemented with appropriate antibiotics: 20 µg/ml kanamycin (Km) or 100 µg/ml ampicillin (Amp). The *Msm* wild-type strain mc^2^155 and its mutant derivatives were grown at 37°C in either (1) Middlebrook 7H9 broth (Difco), supplemented with 0.05% Tween 80, 0.4% glycerol, and 5% albumin-dextrose-catalase (ADC, Sigma-Aldrich) on a shaking platform, or (2) on Middlebrook 7H10 agar, supplemented with 0.4% glycerol and 5% oleic-albumin-dextrose-catalase (OADC, Sigma-Aldrich). When necessary, antibiotics were added at the following concentrations: 20 µg/ml kanamycin, 25 µg/ml hygromycin (Hyg), or 50 µg/ml Zeocin (Zeo). Bacterial growth was monitored by measuring the OD600 at various time points.

### Plasmid and oligonucleotides

All plasmids used in this study are listed in (**Supplementary Table S3)**. The plasmid pJV53 is an *Escherichia*-*Mycobacteria* shuttle plasmid expressing the gp60-61 recombinase under TetO-regulated promoters (Addgene #26904) ^57^. The pJV53-cas12a is a plasmid expressing (1) cas12a under an Acetamide-regulated promoter and (2) gp60-61 recombinase under TetO-regulated promoters (Addgene #158706) ^58^. The pCR-Zeo plasmid was used for cloning and constitutive expression of crRNAs in *Mycobacterium* (Addgene #158709) ^58^. All oligonucleotides were ordered from Eurogentec, Belgium, and listed in (**Supplementary Table S4)**.

### General protocol of DNA manipulation

Plasmid purification was conducted using the GeneJet Plasmid Midiprep or Miniprep Kit (Thermo Fisher Scientific). DNA digestion was carried out with FastDigest restriction enzymes (Thermo Fisher Scientific) unless stated otherwise. PCR reactions were performed using OneTaq® DNA polymerase (New England Biolabs) or DreamTaq DNA polymerase (Thermo Fisher Scientific). PCR product purification was done using the GeneJet PCR Purification Kit (Thermo Fisher Scientific). Sequencing was carried out by Eurofins Genomics, Ebersberg Germany, and DNA synthesis by Twist Bioscience.

### Construction of sgRNA Expression Plasmids

The crRNAs targeting specific regions for mutating a particular gene in *M. smegmatis* were designed using CHOPCHOP ^59^. Custom PAM (Protospacer Adjacent Motif) sequences “NGG” were utilized to identify target sites. Two complementary oligonucleotides containing the target sequence were synthesized and annealed to create a protospacer cassette with BpmI and HindIII overhangs at the 5’ and 3’ ends, respectively. This cassette was then cloned into the pCR-Zeo plasmid for the constitutive expression of crRNAs ^58^.

### Preparation of *M. smegmatis* electro-competent cells

An overnight starter culture (10 ml) of *Msm* wild-type strain mc^2^155 was diluted into 100 ml of Middlebrook 7H9 medium and grown at 37 °C with agitation (150 rpm) to an OD600 of 0.8-1.0. The culture was chilled on ice for 1h and then bacteria were harvested by centrifugation (4000g, 15 min, and 4°C), washed three times with cold 10% glycerol, and finally resuspended in 10% glycerol at 1/500th of the original volume ^60^. The plasmids pJV53 and pJV53-Cas12a were electroporated into *Msm* strain mc2155 and plated on a selective medium containing kanamycin (20 µg/ml). The resulting strains *Msm* (pJV53) and *Msm* (pJV53-Cas12a) were cultured at 37°C until an OD600 of 0.45-0.5, then acetamide (0.2%, w/v) was added to induce Che9c 60-61 recombinases. After 3h of induction, electro-competent cells were prepared as described above for the wild-type strain. All electroporations were carried out at 2.5 kV, 25 μF in 2 mm cuvettes using an Electroporator 2510 from Eppendorf.

### Construction of NucS^Tyr132Ser^, MutM^Glu256Asp^, Fpg2^Tyr50His^, XthA^Trp85Arg^ mutants in *Msm*

The NucS^Tyr132Ser^ allele was constructed using the CRISPR-Cas12a-assisted recombineering system ^58^. First, two complementary oligonucleotides targeting *nucS* (CrNucS(132)_Fs and CrNucS(132)_Rs **(Supplementary Table S4)** were annealed and cloned into pCR-Zeo plasmid between *Bpm*I and *Hind*III restriction sites. The resulting pCR-Zeo-nucS-Tyr132Ser plasmid (100 ng) was then electroporated into *Msm* (pJV53-Cas12a) competent cells together with the ssDNA oligonucleotides carrying the point mutation (500 ng). The pCR-Zeo plasmid contains the temperature-sensitive replication origin of pAL5000 ^61^. The electroporated cells (100 µl) were then added to 1 ml 7H9 broth containing 10 ng/ml anhydrotetracycline (ATc), incubated for 4 h at 30°C at 200 rpm, and finally plated on selective 7H10 agar medium containing kanamycin (20 µg/ml), and zeocin (50 µg/ml). After 6 days of growth at 30°C, several colonies were isolated and picked for PCR and sequencing analysis to confirm the desired recombinants. The MutM^Glu256Asp^, Fpg2^Tyr50His^, and XthA^Trp85Arg^ mutants were obtained by allelic exchange ^57^. To perform and select the allelic replacement, linear DNA substrates were commercially synthesized (Twist Bioscience) as follows: a codon-optimized promoterless zeocin resistance cassette is flanked: i) upstream by 300-500 bp of the target sequence containing the mutation to be introduced into the chromosome and ii) by 300-500 bp of the region downstream of this sequence, to allow homologous recombination (fragment sequences are presented in **Supplementary Table S5)**. *Msm* (pVJ53) competent cells, prepared as described above, were electroporated with 100 ng of DNA substrate and incubated in 1 ml of 7H9 broth with 0.2% of acetamide for 4 h at 37°C at 150 rpm. Recombinant clones were selected after 5 days of growth at 37°C on plates containing 50 μg/ml zeocin. PCR and sequencing were then used to verify the correct allelic exchange and the presence of the appropriate mutation on the chromosome **(Supplementary Table S6)**.

### Analyses of mutation frequencies

*Msm* cultures (50 mL) were grown in 7H10 to OD600nm = 0.7-1.0 and 100 µl of each culture was spread onto 7H10 containing rifampicin (100 µg /ml) or Isoniazid (100 µg/ml) to determine the number of spontaneous rifampicin/Isoniazid resistant mutants. Each of the cultures was serially diluted and 100 µl of the appropriate dilution was plated in triplicate on 7H10 media to determine the total number of cells (CFU/mL-Number of colonies*dilution factor) / volume of culture plate). The frequency was calculated as the ratio of Rifampicin^R^/Isoniazid^R^ mutants to the total number of cells. Statistical analysis (one**-**way ANOVA) was performed using n = 6 for each biological experiment in RStudio ^49^.

